# Association of environmental markers with childhood type 1 diabetes mellitus revealed by a long questionnaire on early life exposures and lifestyle in a case-control study

**DOI:** 10.1101/063438

**Authors:** F Balazard, S Le Fur, S Valtat, Isis-Diab collaborative group, AJ Valleron, P Bougnères

## Abstract

**Background:** The incidence of childhood type 1 diabetes (T1D) incidence is rising in many countries, supposedly because of changing environmental factors, which are yet largely unknown.

**Purpose:** To unravel environmental markers associated with T1D. Methods: Cases were children with T1D from the French Isis-Diab cohort. Controls were schoolmates or friends of the patients. Parents were asked to fill a 845-item questionnaire investigating the child’s environment before diagnosis. The analysis took into account the matching between cases and controls. A second analysis used propensity score methods.

**Results:** We found a negative association of several lifestyle variables, gastroenteritis episodes, dental hygiene, hazelnut cocoa spread consumption, wasp and bee stings with T1D, consumption of vegetables from a farm and death of a pet by old age.

**Conclusions:** The found statistical association of new environmental markers with T1D calls for replication in other cohorts and investigation of new environmental areas.

## Background

The current rise in T1D incidence [1] is attributed to environmental causes to which genetically predisposed children are increasingly exposed, but epidemiology has delivered more questions than robust answers. Dissecting the environment is a daunting task, with paramount difficulties for extracting relevant information from multiple known and unknown exposures occurring during childhood. The fact that childhood T1D occurs early in life allows restraining the environmental analysis to the few years encompassing intrauterine life, infancy and childhood. A classical way of doing this is using retrospective questionnaires, but the questions are necessarily limited to selected areas of child life and answers may be biased by parental recall. Environmental comparison between cases and controls can also be prospective. To achieve this given the low prevalence of T1D, it is necessary to study a genetically at risk population, for example positivity for HLA screening in the TEDDY study [2]. Another way of avoiding recall-related problems is to use registries [3]. However, registries are more limited in their scope than a questionnaire. Another difficulty inherent to any environmental approach is that participants are not aware of many exposures. Collecting biological samples to characterize the “exposome” [4] of T1D children also has several drawbacks, since blood parameters may be modified as a consequence of T1D not as a causal component, and are confined to the only environmental parameters that leave a long living trace in patient’s blood, i.e. a minority of exposures.

Over the recent years, suspicion has almost exclusively focused on infectious agents and nutrition in the early years of life [5–7]. Enteroviruses have been the subject of numerous studies and have remained the most often suspected environmental contributors to T1D [8, 9]. In contrast, infections have been considered as protective from T1D according to the hygiene hypothesis, which postulates that the increase in autoimmune T1D could be due to the decrease of early infections [10, 11] or lack of parasites [12][12] This has been shown in the isogenic NOD mice model [11, 13], but epidemiological evidence in humans, who are exposed to different infectious agents and have a wide genetic variation, is still pending. Studies attempting to relate infectious episodes with T1D have yielded contrasted results [14]. Respiratory infections in the first year of life have been shown to increase the risk of seroconversion to islet autoimmunity (IA) in the BABYDIET cohort and in the MIDIA study [15, 16]. On the other hand, they were not associated with T1D in the DAISY cohort [17]. Gastrointestinal illnesses at precise periods were associated with higher risk of IA in the same study. More recently, the gut microbiome has been investigated in search of a bacterial composition that could be associated with T1D [18].

Nutrition has been the other focus of environmental research for T1D. Overfeeding and the ensuing increase of beta cell functional activity for producing more insulin has been suspected to favor autoimmunity towards the beta cells (the overload hypothesis) [19]. Meta-analyses have found that early weight gain [20] or obesity [21] showed a modest association with T1D. Vitamin D supplementation studied through questionnaires has been suggested to protect from T1D [22], but this has not been confirmed when 25-hydroxyvitamin D levels in plasma were studied [23]. Since vitamin D supplementation of infants is generalized in French infants since the 70s, vitamin D deficiency is not likely to be a driver of increasing T1D incidence.

Several dietary interventions have attempted to prevent T1D. TRIGR tested whether substitution of cow’s milk by casein hydrolysate formula affects the occurrence of IA or progression to T1D [24]. No significant difference has been observed between the two groups for the appearance of IA [25]. Result for the second primary end-point will only be available in 2017. The possibility that exclusive breast-feeding or late introduction to cow’s milk is associated with a modest protection is supported by a meta-analysis of observational studies [26]. A few other nutrients have been studied. An older age at first introduction to gluten showed no protective effect in the BABYDIET study [27]. Omega-3 fatty acids seemed to be associated with a slightly reduced risk of islet autoimmunity in the DAISY cohort [28] but the pilot study that was then performed did not show significant protection [29]. Nicotinamide did not modify the progression to T1D in children with IA in the ENDIT trial [30]. Other prevention trials are underway [31]. Early nutrition is a favorable field of investigation through randomized trials since a vast number of factors can be manipulated experimentally.

The BABYDIAB and the DAISY cohort have found that IA often appears in the first years of life preceding clinical diagnosis T1D by several months or years [32, 33], which stress a potential predisposing role for early environmental exposures. This has inspired our approach for screening early life events that could be associated with environmental differences between cases and controls, including a number of infectious and nutritional exposures that can be reliably recalled by parents.

Our study is a tentative and still limited step for moving environmental research from hypothesis-driven to more data-driven approaches. A comparable move has occurred in the 90s when genetic research has switched its candidate gene approach of complex diseases, notably T1D, to interrogate the complete genome variation blindly with genome wide association studies (GWAS) with the aim of unraveling disease markers [34] that could secondarily lead to true genetic causation [35, 36]. Environment wide association studies (EWAS) [37] or exposome association studies [4] will likely allow researchers to investigate children environment on a vast scale without making a priori hypotheses. Such approaches will remain limited because a myriad of environmental markers will escape investigation, while genomic variation is finite. In this respect, our current 845-item questionnaire can only be viewed as a preliminary proof-of-concept approach for scanning the environment of a child. It is indeed limited by the number of questions that have been selected to describe this environment, by the recall errors that could be made by the parents of the cases and controls, and by the number of participants who agree to spend two hours filling a complex questionnaire. False positivity is an expected weakness of this approach, but careful statistical analysis can provide a list of environmental markers for which false discoveries are controlled.

## Methods

### Questionnaire

The questionnaire was built by a group of academics composed of obstetricians and pediatricians specialized in pediatric infectious diseases, nutrition, and lifestyle. Their task was to define the environment of pregnant women, neonates, infants and young children, by enumerating all aspects that they thought a mother will likely be able to recall years later. A group of mothers of young children (living in urban or rural environment) were also asked to participate. A first questionnaire of nearly 1,000 questions was built and tested across 100 young mothers. Only questions that could be answered rapidly were kept, because we considered that the speed of the answers would favor spontaneity and minimize recall errors and bias. The questionnaire was also tested in 30 mothers of young children with recently diagnosed T1D and 30 mothers of children who had declared T1D five to ten years before. Only questions that had a comparable recall score in the two groups of mothers were kept in an effort to eliminate questions that could not be easily recalled. The final questionnaire contained 576 main questions and 845 items when counting sub-questions about the environment including 90 questions about pregnancy, 25 concerning the delivery and early post-natal life, 20 about early childhood, 75 on the subject’s medical life, 60 on nutrition, 40 on housing, 30 on daycare, 30 on leisure and trips, 80 on contact with animals and 60 on family members’ environment. Depending on mothers, the time to fill the questionnaire ranged form 90 to 120 minutes. A PDF version of it in French is available as supplementary file 2.

### Data collection

The Isis-Diab cohort is a large multi-centric cohort of T1D patients in France which recruitment started in 2007. After march 2010, three copies of the questionnaire were sent to the parents of 6618 T1D patients enrolled in the cohort during the month following their inclusion in the study. Parents were asked to fill the questionnaire regarding the exposures and events having taken place in their child’s life before the clinical onset of T1D. They were also asked to enroll as controls two of their friends having an unaffected child of the same age. The 6144 parents having provided a phone number were contacted once during the week following the questionnaire sending. If the questionnaire was not returned within 3 months, parents received a reminder by mail.

1769 cases (i.e. 27% of the patients to which a questionnaire was sent) and 1085 controls returned the questionnaire. 241 cases provided two controls, 451 cases provided one control, and 1077 cases provided no control. 152 controls were not associated to a case that returned his questionnaire. All the questionnaires completed by patients and controls were seized by a private provider (numerical input for all the « checkbox » responses, and dual manual entry for handwritten responses).

Patients living in areas with higher economic deprivation were less likely to respond [31].

The questionnaire investigated the period preceding diagnosis of the disease. Matched controls were asked to fill the questionnaire with respect to the age at which the patient had been diagnosed. We will refer to this age as the reference age.

### Pre-analysis treatment

A computerized treatment was designed to code categorical questions into binary variables and to allow analysis of subquestions. After the pre-analysis treatment, 845 variables were available for analysis.

In order for effect sizes to be on a similar scale even though we have binary questions and ordinal questions with up to 5 different levels. For example, consumption of cola drinks frequency was quantified on five levels from never to several times a day. All variables were scaled to be between 0 and 1. In this way, the effect size for ordinal variables corresponds intuitively to the odds ratio between the two extreme responses. The encoding of the variables were modified so that the directionality of the effects be intuitive. A description of the 845 variables is available in supplementary file 3.

### Exclusions

We excluded from the analysis the questionnaires where more than 50% of the questions were left unanswered.

As our questionnaire was designed to quantify a child’s environment, we included only participants whose reference age was between 0.5 years and 15.5 years. To minimize recall errors, we excluded participants for whom the delay between diagnosis and questionnaire reception was greater than 10 years.

We used primary school attendance as another marker of the quality of recall: we excluded participants who reported that their child attended primary school before 5.5 years. In the supplementary material, we use a questionnaire-based prediction model for age to justify this exclusion. Using the same prediction model, we consider a second exclusion of outliers for predicted age in the supplementary material. We report which results are significantly affected by this further exclusion.

For the first analysis, we excluded participants without matched counterparts, i.e. patients with no matched control or controls with no matched patient. The matched analysis then compared 469 patients with 624 matched controls.

We also performed a propensity analysis without using the matching. We only excluded participants with no available postal code or parents’ profession as these variables were used to control for bias. This resulted in a sample of 1151 patients and 689 controls. The processes of exclusion and sample definition are summarized in figure 1.

**Fig. 1.**
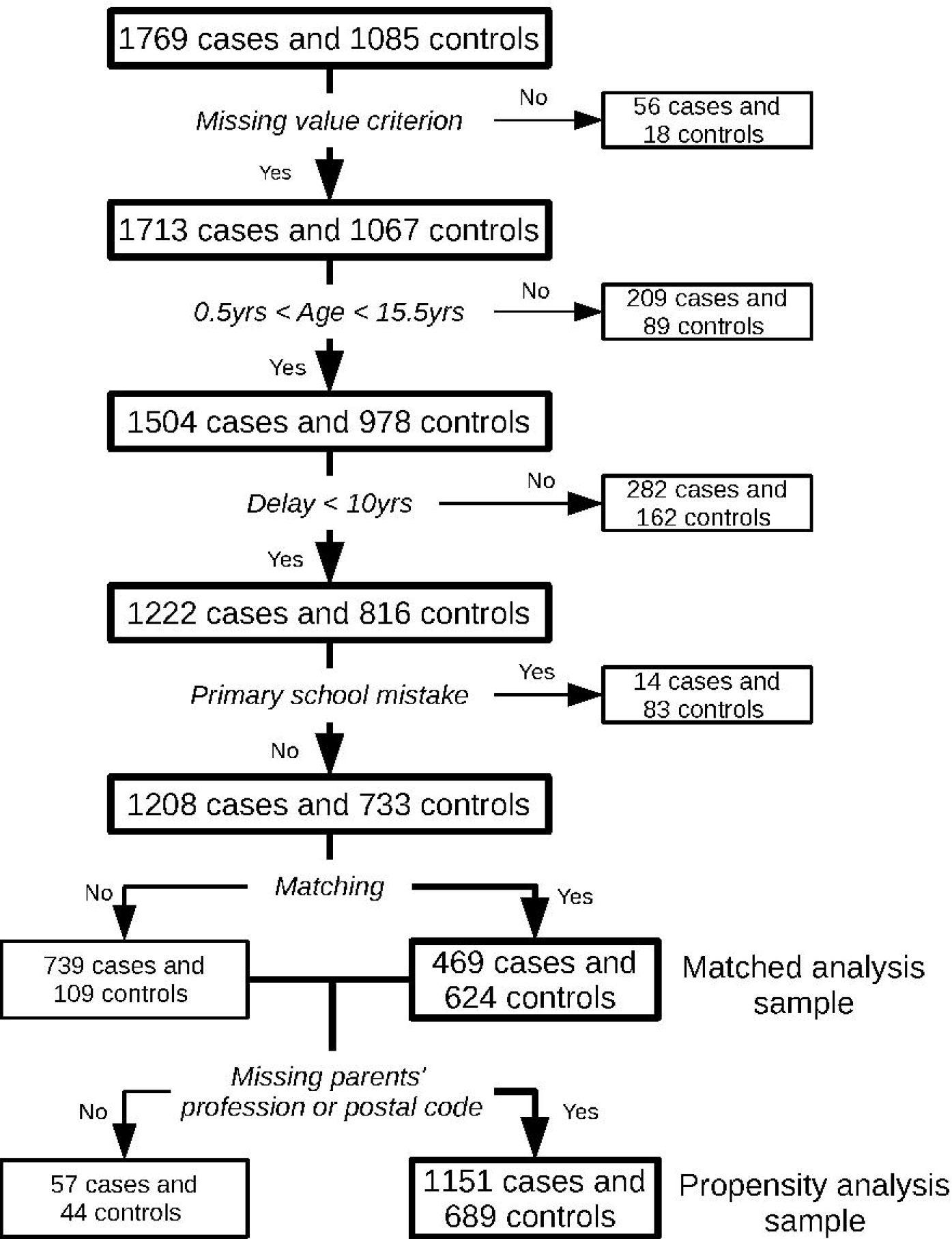
Flowchart of the samples definition. Missing value criterion is verified if at least half the questionnaire was filled. Delay refers to the time between diagnosis and questionnaire reception. Participants have made the primary school mistake if they answered that they went to primary school even though their reference age is smaller than 5.5 years. The two samples on which analyses were performed are in the bottom right corner.

## Analytical procedures

### Matched analysis

We used methods that take matching into account and allow for variable size of the matched strata: either one patient and one control or one patient and two controls. For questions with binary responses, we performed Cochran-Mantel-Haenszel tests and for ordinal responses, we performed conditional logistic regression [38]. In both cases, we used the strata defined by the matching and the disease status as outcome. To avoid convergence problems, we excluded variables with a standard deviation smaller than 0.1.

### Propensity analysis

In this second analysis, we used stratification on the propensity score [39] to control for bias. Propensity score methods allow to control for bias by comparing participants with a similar probability of treatment (here the response to a question) given the covariates defined below.

Random Forests [40 p587-604] is a popular machine learning algorithm praised for its state-of-the-art predictive performance. Furthermore, it provides a reliable prediction on the training set called out-of-bag estimate which is not prone to overfitting. We used the randomForest package in R [41]. We trained a Random Forests regression to predict the treatment status using as predictors reference age, socio-economic status, urban/rural environment and study center. We then defined the propensity score as the out-of-bag estimate of the random forest. We then stratified our sample in 10 strata according to deciles of the propensity score and performed a Cochran-Mantel-Haenszel test (respectively a conditional logistic regression) between the question of interest if it was binary (respectively if it was ordinal) and disease status. We again excluded variables whose standard deviation was smaller than 0.1.

### Covariate description

The following covariates were used to define the propensity score for the propensity analysis:

#### Age

The reference age was written on the first page of the questionnaire as an integer number of years that corresponds to a rounding of the patient’s age at diagnostic. In both analyses, we used non-rounded patient’s age at diagnostic for both the patient and his matched controls.

#### Socio-economic status

Socio-economic status was assessed using the hand-written professions of parents. It was encoded as an ordinal variable taking value 0, 1 or 2 where 0 corresponds to blue-collar workers, 1 to intermediate professions and 2 to upper class. Among the 1840 participants of the propensity analysis, 837 were classified as 0, 725 as 1, and 278 as 2.

#### Urban/farm environment

Using the postal code of the participants obtained through the questionnaire, two variables defined at the level of the patient’s “commune” (town) of residence were used to quantify whether the participants lived in an urban or rural area. Those variables are the urban units index (as a code reflecting the size of the commune’s urban area) and the percentage of farmers in the active population. Those two variables came from anonymous public databases (French Quetelet Network (http://www.reseau-quetelet.cnrs.fr), via the Centre Maurice Halbwachs-Archives de Données Issues de la Statistique Publique (http://www.cmh.greco.ens.fr/adisp.php)) and were dated in 2007 (census closest to the date that patients started to receive the environmental questionnaire). Environment was also controlled by the recruitment center e.g. the hospital or pediatric endocrinology practice that recruited the patient: each center with more than 30 participants was coded as a distinct binary variable.

### Correction for multiple tests

To control for multiple testing, we used the Bonferroni correction which allows to control the family-wise error rate at 5%. For the matched analysis, as we consider that it is of better quality than the propensity analysis, we also considered the more lenient false discovery rate [42] for a level of 5%.

We report the list of variables that passes both the FDR threshold for the matched analysis and the Bonferroni threshold for the propensity analysis. This provides better control over false positives than considering only one of the two thresholds. We also report results for variables associated with T1D in the literature.

### Code

Code used for analysis as well as pre-analysis treatment is available on github: https://github.com/FelBalazard/ISIS_DIAB

### Results

**Table 1:** Characteristics of cases and controls in the two samples. Age is the reference age. Delay is the time between the diagnosis date and the questionnaire reception. The values displayed are the median value and the first and third quartile between parentheses.

|  | Matched analysis sample |  | Propensity analysis sample |  |
| --- | --- | --- | --- | --- |
|  | Cases | Controls | Cases | Controls |
| Number | 469 | 624 | 1151 | 689 |
| Age (years) | 7.5 (4.2;10.5) | 7.7 (4.6;10.5) | 7.6 (4.2;10.6) | 7.8 (4.6;10.5) |
| Delay (years) | 3.0 (1.0;5.2) | 2.9 (1.1;5.2) | 2.9 (0.9;5.6) | 3.0 (1.1;5.4) |
| Missing data (%) | 4.4 (2.7;6.8) | 3.9 (2.5;6.0) | 4.9 (3.1;7.5) | 3.8 (2.4;5.9) |

### Matched analysis

For convenience, the variables have been labeled in the figures. Correspondence between labels and precise description of variables are available in supplementary file 3.

Figure 2 presents a volcano plot where both the effect size and the significance of answers to each question are displayed. We also display in blue the Bonferroni-Holm threshold for multiple testing, this means that we control the family-wise error rate at 5% for the list of variable over the blue line. The more lenient threshold for a false discovery rate of 5% is displayed in red. Questions that pass this threshold are labeled in the plot. Exact sample size, p-value, estimate and confidence interval for each variable are available as supplementary file 3.

Three questions showed that cases more often had a relative with T1D and are excluded from the plots and discussion.

**Fig. 2.**
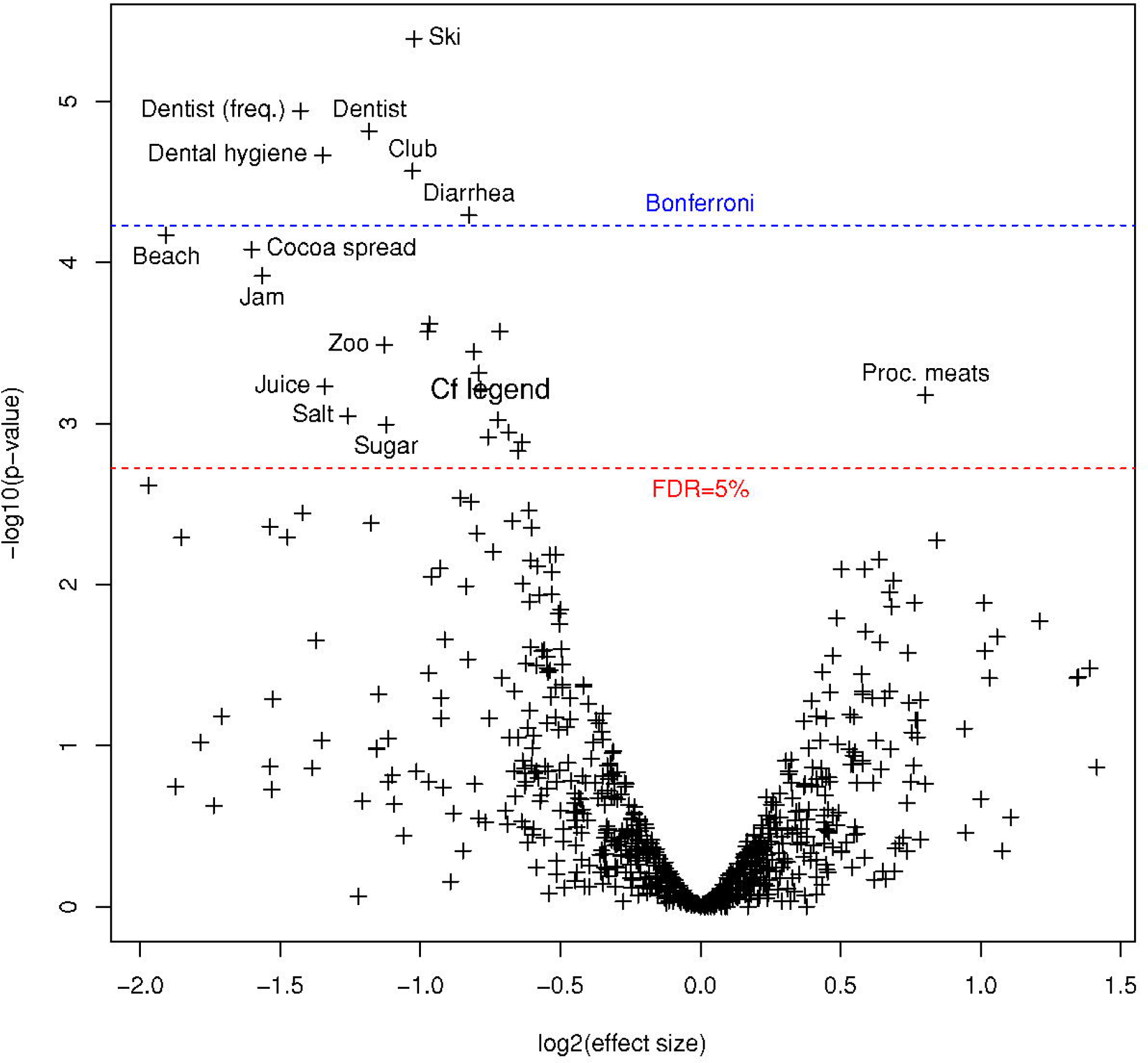
Volcano plot for the matched analysis. The x-axis shows the effect size with protective factors on the left and risk factors on the right. The y-axis indicates the significance. The higher line indicates the Bonferroni threshold while the lower line shows the more lenient threshold for 5% of false discovery rate. The unlabeled variables above the FDR threshold are from most significant to least: week-ends with other children, taste for sugar as a baby, death of a pet from old age, vegetables from farm, home-made delicatessen, stings (mainly wasps and bees), siblings before birth, friend’s pool, plane, fresh exotic fruits, vegetables from a rural market during pregnancy.

### Propensity analysis

Results are also available in supplementary file 3. They are shown in figure 3.

**Fig. 3.**
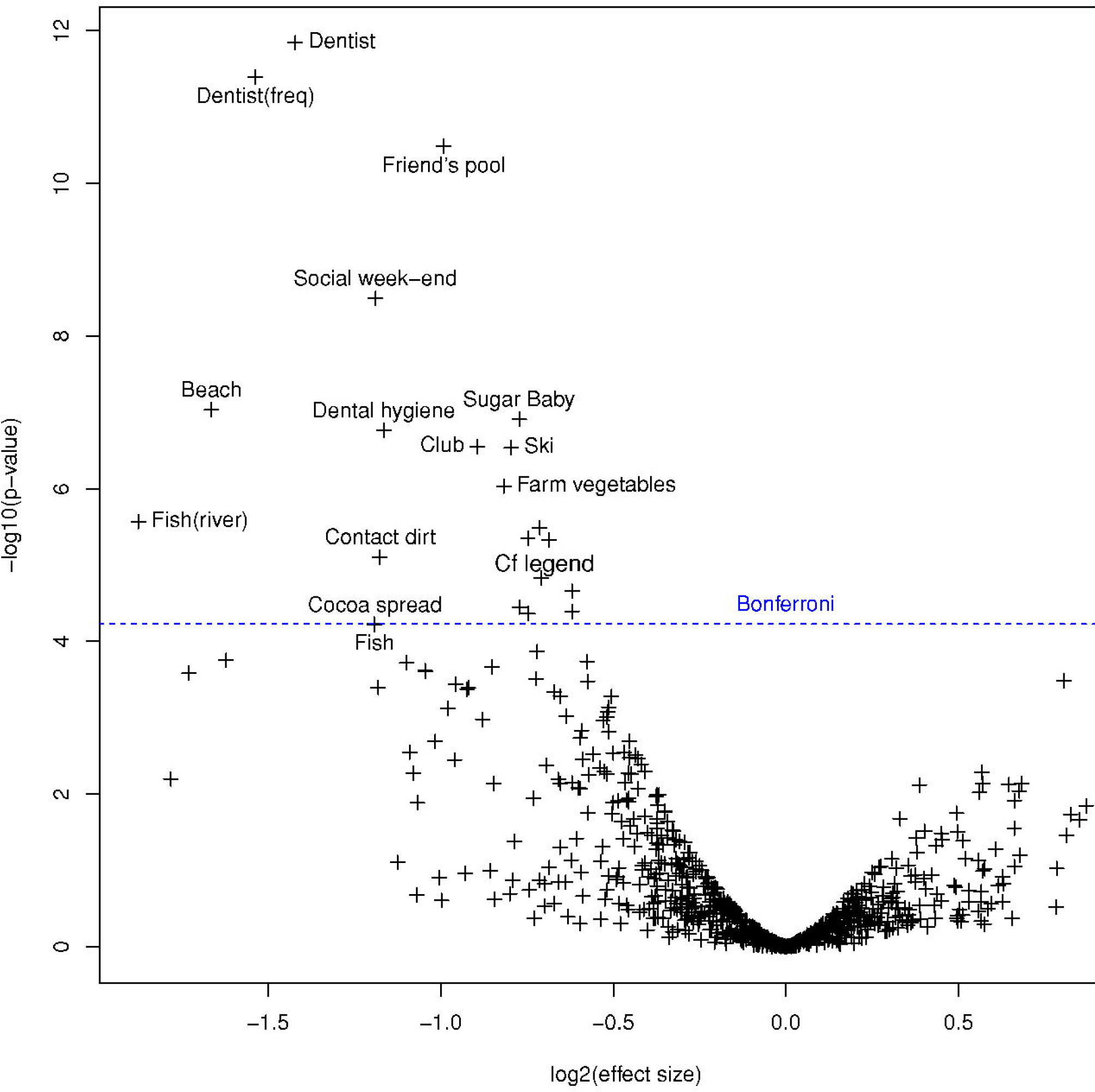
Volcano plot for the propensity analysis. The x-axis shows the effect size with protective factors on the left and risk factors on the right. The y-axis indicates the significance. The horizontal line indicates the Bonferroni threshold. The unlabeled variables above the threshold are from most significant to least: fruits from a farm or a family garden during childhood, stings, diarrhea, diarrhea during winter, contact with cats in the neighborhood, pet shop, swimming pool during pregnancy and death of a pet of old age.

### Comparison

**Fig. 4.**
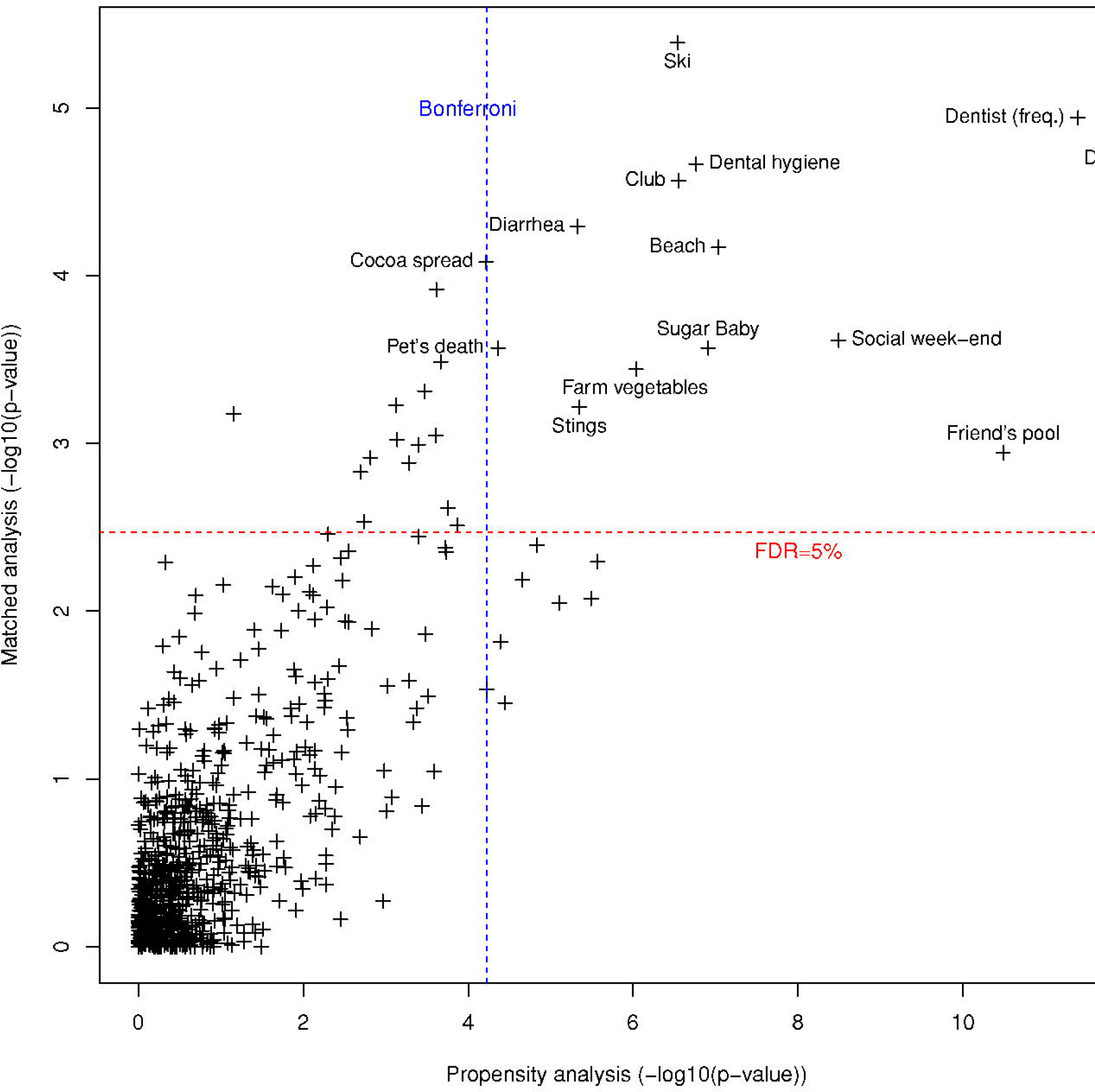
Comparison of the results of the two analysis. -log10(p-value) of the two analysis plotted against each other. The most associated variables in both analysis are in the top right corner. The Bonferroni threshold for the propensity analysis is the vertical line. The false discovery rate threshold for the matched analysis is the horizontal line. A more lenient statistical control is used for the matched analysis as it is less prone to bias. All variables passing both thresholds are labeled.

The result of the two analyses are summarized in figure 4.

Social variables and markers of outdoor life are negatively associated with T1D: club attendance, playing with friends during the week-end, going to the pool at a friend’s house, winter sports and going often to the beach. Going often to the beach was sensitive to the age-related exclusion considered in the supplementary material. Club attendance was also partially affected.

Patients had less gastroenteritis before T1D diagnosis.

Hazelnut cocoa spread consumption and sweet eating as a baby were both negatively associated with T1D.

Three variables were closely connected to dental hygiene. The variable “dental hygiene” is an ordinal variable quantifying the frequency at which the participants brush their teeth. The two variables “dentist” and “dentist (freq.)” are a binary and an ordinal variable quantifying the number of dentist visit attended by the participant. Future T1D patients attended the dentist less and brushed their teeth less as well. The association for dentist attendance was very sensitive to the further exclusion considered in the supplementary material. Dental hygiene was also partially affected. The patients reported having been stung less than controls. “Stings” refers to the question: Was the subject stung by an animal who left a clear spot (red spot, painful or not)? with four propositions for the responsible animal: a wasp, a bee, another insect or a fish. Mosquitoes, spiders and ticks were the subjects of separate questions. Wasp and bee stings were the most common stings.

Patients less often had the experience of having a pet die of old age.
Patients ate less vegetables coming from a farm or a family garden.

### Factors studied in the literature

We compared the results of our study with the few risk factors that have been suspected to be associated with T1D in the studies cited in the introduction.

Breastfeeding was investigated by two questions in the questionnaire: whether the subject had been breastfed at all and the duration of exclusive breastfeeding. In the matched analysis, neither questions were significant at the nominal level but in the propensity analysis, the duration of exclusive breast-feeding was found to be highly protective. Any breastfeeding was also protective with nominal significance.

Lower respiratory infections were not associated with risk of T1D in our analyses.

Vitamin D supplementation for the mother after birth was not associated with T1D in either analysis.

**Table 2.**
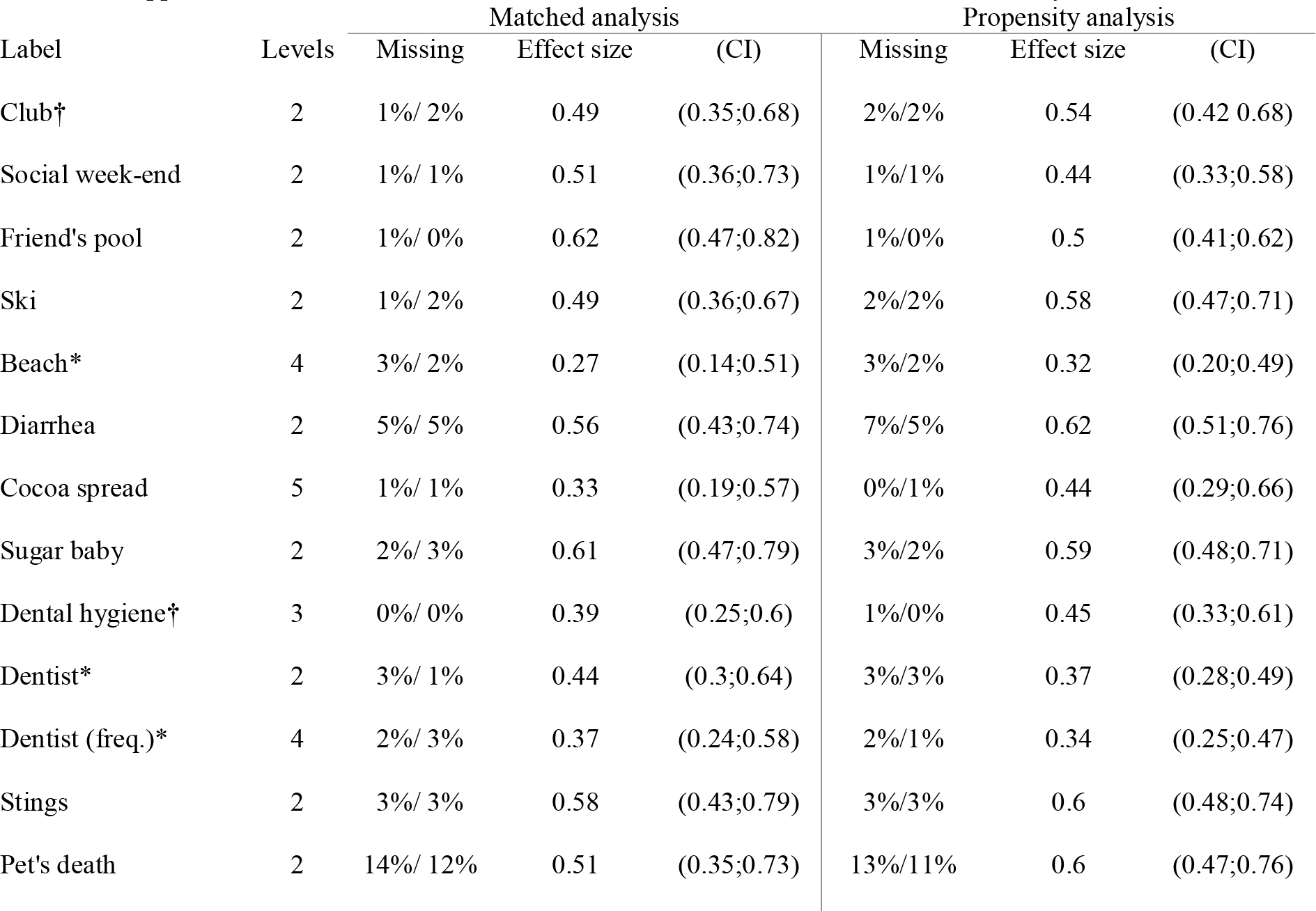
Effect sizes for significant variables and pending risk factors. Effect sizes are odd ratios for binary variables and correspond to odd ratio between extreme responses for ordinal variables. Percentage of missing data are split between patients and controls. Factors from the literature are at the end of the table. *: variables affected by further age-related exclusion. †: variables affected by the further exclusion for the propensity analysis only.

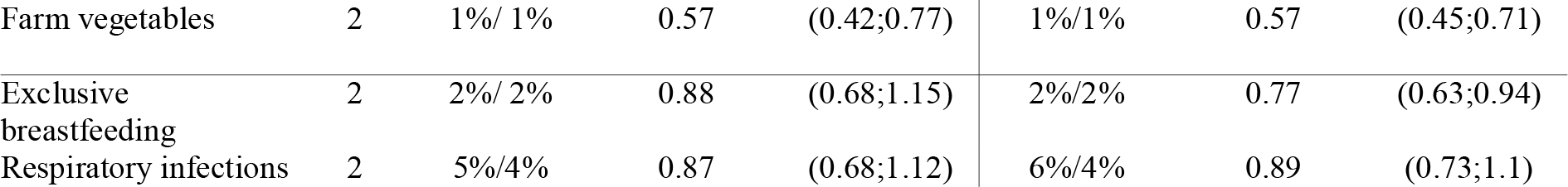

## Discussion

While our statistical analysis indicates that playing with friends during week-ends or going to the pool at a friend’s house and experience of winter sports were all negatively associated to childhood T1D, we have not attempted to interpret these protective associations.

We also found a negative association of gastroenteritis and T1D. Gut microbiology is an area highlighted by this observation. Sub-questions regarding gastroenteritis reveal that the negative association holds for diarrhea during winter and in the context of familial diarrhea. The results of the DAISY study suggested a more complex relationship with gastroenteritis [17].

As sugar consumption is strongly present as a nutritional caveat in the minds of parents having a child with T1D, we suspected that the negative association between “appetite for sugar as a baby” and T1D could be due to recall bias. However, with respect to a possible recall bias, sugary products such as cola drinks or chocolate show no association with T1D. This gives credibility to the found negative association for hazelnut cocoa spread. Furthermore, hazelnut cocoa spread remains significant after adjustment for appetite for sugar as a baby: in the matched sample, fitting a conditional logistic regression to both variables gives an estimate for cocoa spread of 0.36 (0.20,0.64) instead of 0.33 (0.19,0.57), meaning that the result for cocoa spread is not affected by recall bias. Hazelnut cocoa spread contains a large proportion of palm oil thus a high content of tocotrienol. In murine models, tocotrienol was shown to affect NLPR3 and NF-κB [43, 44], which may play a role in T1D pathogenesis [45, 46].

We found that items related with dental hygiene, such as frequency of teeth brushing and dentist attendance, were negatively associated with T1D although they were sensitive to a further exclusion. Again, we have not attempted to interpret this protective association in our current state of knowledge.

Wasp and bee stings also showed a significant association with T1D, but the meaning of this observation remains to be found.

Death of pet by old age was negatively associated with T1D. This was a subquestion of death of a pet which was nominally significant in both analysis. Another subquestion, death of a cat, was also associated. We offer no interpretation.

Eating vegetables from a farm or a family garden was negatively associated with T1D. The analogue question for fruits passed the Bonferroni threshold in the propensity analysis and was also nominally significant in the matched analysis. These associations might be connected to contact with dirt which was also significant in the propensity analysis and nominally significant in the matched analysis. Again, we offer no interpretation.

In conclusion, while many exposures and events have remained out of reach of our questionnaire because they were not detectable or escaped parental memory, the novel protective associations that were found cannot be entirely false positive findings. They may open new areas of investigation for T1D environmental research and should not be dismissed more than yet biologically inexplicable SNP associations generated by GWAS. However they will only be of interest if they can be confirmed in other childhood T1D cohorts.

## Acknowledgement

We thank Gérard Biau for his oversight. We thank Alain Fourreau, Adeline Guégan, Gaël Leprun and Valérie Jauffret for mailing the questionnaires and entering the responses. We thank the participants and their parents for their time.

## Compliance with ethical standards

Patients were included in the study according to the French bioethics law with families being carefully informed and having signed a detailed informed consent agreed by CPP (number DC-2008-693; NI 2620, Comité de Protection des Personnes). Clinical-Trial.gov identifier: NCT02212522.

## References

1. The DIAMOND Project Group (2006) Incidence and trends of childhood Type 1 diabetes worldwide 1990-1999. Diabet Med 23:857–866. doi:10.1111/j.1464-5491.2006.01925.x

2. Hagopian WA, Lernmark A, Rewers MJ, et al (2006) TEDDY--The Environmental Determinants of Diabetes in the Young: an observational clinical trial. Ann N Y Acad Sci 1079:320–326 doi:10.1196/annals.1375.049

3 D’Angeli MA, Merzon E, Valbuena LF, et al (2010) Environmental factors associated with childhood-onset type 1 diabetes mellitus: an exploration of the hygiene and overload hypotheses. Arch Pediatr Adolesc Med 164:732–738. doi:10.1001/archpediatrics.2010.115

4 Rappaport SM, Barupal DK, Wishart D, et al (2014) The blood exposome and its role in discovering causes of disease. Environ Health Perspect 122:769–774 doi:10.1289/ehp.1308015

5 Egro FM (2013)Why is type 1 diabetes increasing? J Mol Endocrinol 51:R1–13. doi:10.1111/j.1464-5491.2006.01925.x 10.1530/JME-13-0067

6 Knip M, Simell O, (2012) Environmental Triggers of Type 1 Diabetes. Cold Spring Harb Perspect Med. doi:10.1101/cshperspect.a007690

7 Forlenza GP, Rewers M (2011) The epidemic of type 1 diabetes: what is it telling us? Curr Opin Endocrinol Diabetes Obes 18:248–251 doi:10.1097/MED.0b013e32834872ce

8 Green J, Casabonne D, Newton R (2004) Coxsackie B virus serology and Type 1 diabetes mellitus: a systematic review of published case-control studies. Diabet Med J Br Diabet Assoc 21:507–514 doi:10.1111/j.1464-5491.2004.01182.x

9 Yeung W-CG, Rawlinson WD, Craig ME, (2011) Enterovirus infection and type 1 diabetes mellitus: systematic review and meta-analysis of observational molecular studies. BMJ 342:d35.

10 Chapman NM, Coppieters K, von Herrath M, Tracy S (2012) The microbiology of human hygiene and its impact on type 1 diabetes. Islets 4:253–261 doi:10.4161/isl.21570

11 Bach J-F (2002) The Effect of Infections on Susceptibility to Autoimmune and Allergic Diseases. N Engl J Med 347:911–920 doi:10.1056/NEJMra020100

12 Gale E a. M (2002) A missing link in the hygiene hypothesis? Diabetologia 45:588–594 doi:10.1007/s00125-002-0801-1

13 Like AA, Guberski DL, Butler L (1991) Influence of environmental viral agents on frequency and tempo of diabetes mellitus in BB/Wor rats. Diabetes 40:259–262

14 Schneider DA, Herrath MG (2014) Potential viral pathogenic mechanism in human type 1 diabetes. Diabetologia 57:2009–2018 doi:10.1007/s00125-014-3340-7

15 Beyerlein A, Wehweck F, Ziegler A-G, Pflueger M (2013) Respiratory infections in early life and the development of islet autoimmunity in children at increased type 1 diabetes risk: evidence from the BABYDIET study. JAMA Pediatr 167:800–807 doi:10.1001/jamapediatrics.2013.158

16 Rasmussen T, Witsø E, Tapia G, et al (2011) Self-reported lower respiratory tract infections and development of islet autoimmunity in children with the type 1 diabetes high-risk HLA genotype: the MIDIA study. Diabetes Metab Res Rev 27:834–837 doi:10.1002/dmrr.1258

17 Snell-Bergeon JK, Smith J, Dong F et al (2012) Early childhood infections and the risk of islet autoimmunity: the Diabetes Autoimmunity Study in the Young (DAISY). Diabetes Care 35:2553–2558 doi:10.2337/dc12-0423

18 Vatanen T, Kostic AD, d’Hennezel E, et al (2016) Variation in Microbiome LPS Immunogenicity Contributes to Autoimmunity in Humans. Cell 165:842–853 doi:10.1016/j.cell.2016.04.007

19 Dahlquist G (2006) Can we slow the rising incidence of childhood-onset autoimmune diabetes? The overload hypothesis. Diabetologia 49:20–24 doi:10.1007/s00125-005-0076-4

20 Harder T, Roepke K, Diller N et al (2009) Birth weight, early weight gain, and subsequent risk of type 1 diabetes: systematic review and meta-analysis. Am J Epidemiol 169:1428–1436 doi:10.1093/aje/kwp065

21 Verbeeten KC, Elks CE, Daneman D, Ong KK (2011) Association between childhood obesity and subsequent Type 1 diabetes: a systematic review and meta-analysis. Diabet Med 28:10–18 doi:10.1111/j.1464-5491.2010.03160.x

22 Dong J-Y, Zhang W, Chen JJ et al (2013) Vitamin D Intake and Risk of Type 1 Diabetes: A Meta-Analysis of Observational Studies. Nutrients 5:3551–3562 doi:10.3390/nu5093551

23 Simpson M, Brady H, Yin X, et al (2011) No association of vitamin D intake or 25-hydroxyvitamin D levels in childhood with risk of islet autoimmunity and type 1 diabetes: the Diabetes Autoimmunity Study in the Young (DAISY). Diabetologia 54:2779–2788 doi:10.1007/s00125-011-2278-2

24 TRIGR Study Group, Akerblom HK, Krischer J, et al (2011) The Trial to Reduce IDDM in the Genetically at Risk (TRIGR) study: recruitment, intervention and follow-up. Diabetologia 54:627–633 doi:10.1007/s00125-010-1964-9

25 Knip M, Åkerblom HK, Becker D, et al (2014) Hydrolyzed infant formula and early β-cell autoimmunity: A randomized clinical trial. JAMA 311:2279–2287 doi:10.1001/jama.2014.5610

26 Cardwell CR, Stene LC, Ludvigsson J, et al (2012) Breast-Feeding and Childhood-Onset Type 1 Diabetes A pooled analysis of individual participant data from 43 observational studies. Diabetes Care DC_120438. doi:10.2337/dc12-0438

27 Hummel S, , Pflüger M, Hummel M, et al (2011) Primary dietary intervention study to reduce the risk of islet autoimmunity in children at increased risk for type 1 diabetes: the BABYDIET study. Diabetes Care 34:1301–1305. doi:10.2337/dc10-2456

28 Norris JM, Yin X, Lamb MM et al (2007) Omega-3 polyunsaturated fatty acid intake and islet autoimmunity in children at increased risk for type 1 diabetes. JAMA 298:1420–1428. doi:10.1001/jama.298.12.1420

29 Chase HP, Boulware D, Rodriguez H, et al (2015) Effect of docosahexaenoic acid supplementation on inflammatory cytokine levels in infants at high genetic risk for type 1 diabetes. Pediatr Diabetes 16:271–279 doi:10.1111/pedi. 12170

30 Gale E a. M, Bingley PJ, Emmett CL et al (2004) European Nicotinamide Diabetes Intervention Trial (ENDIT): a randomised controlled trial of intervention before the onset of type 1 diabetes. Lancet Lond Engl 363:925–931 doi:10.1016/S0140-6736(04)15786-3

31 Skyler JS (2013) Primary and secondary prevention of Type 1 diabetes. Diabet Med 30:161–169 doi:10.1111/dme.12100

32 Barker JM, Barriga KJ, Yu L, et al (2004) Prediction of Autoantibody Positivity and Progression to Type 1 Diabetes: Diabetes Autoimmunity Study in the Young (DAISY). J Clin Endocrinol Metab 89:3896–3902 doi:10.1210/jc.2003-031887

33 Ziegler AG, Hummel M, Schenker M, Bonifacio E (1999) Autoantibody appearance and risk for development of childhood diabetes in offspring of parents with type 1 diabetes: the 2-year analysis of the German BABYDIAB Study. Diabetes 48:460–468 doi:10.2337/diabetes.48.3.460

34 Welter D, MacArthur J, Morales J et al (2014) The NHGRI GWAS Catalog, a curated resource of SNP-trait associations. Nucleic Acids Res 42:D1001–D1006. doi:10.1093/nar/gkt1229

35 Farh KK-H, Marson A, Zhu J et al (2015) Genetic and epigenetic fine mapping of causal autoimmune disease variants. Nature 518:337–343 doi:10.1038/nature13835

36 Stamatoyannopoulos J (2016) Connecting the regulatory genome. Nat Genet 48:479–480 doi:10.1038/ng.3553

37 Patel CJ, Bhattacharya J, Butte AJ (2010) An Environment-Wide Association Study (EWAS) on Type 2 Diabetes Mellitus. PLOS ONE 5:e10746. doi:10.1371/journal.pone.0010746

38 Breslow NE, Day NE (1980) Statistical methods in cancer research. Volume I-The analysis of case-control studies IARC Sci Publ 5–338.

39 Austin PC (2011) An Introduction to Propensity Score Methods for Reducing the Effects of Confounding in Observational Studies. Multivar Behav Res 46:399–424 doi:10.1080/00273171.2011.568786

40 Hastie T, Tibshirani R, Friedman J (2009) The Elements of Statistical Learning. Springer New York, New York, NY

41 Liaw A, Wiener M (2002) Classification and Regression by randomForest. R News 2:18–22

42 Benjamini Y, Hochberg Y (1995) Controlling the false discovery rate: a practical and powerful approach to multiple testing. J R Stat Soc Ser B Methodol 289–300.

43 Kim Y, Wang W, Okla M et al (2016) Suppression of NLRP3 inflammasome by γ-tocotrienol ameliorates type 2 diabetes. J Lipid Res 57:66–76 doi:10.1194/jlr.M062828

44 Kuhad A, Bishnoi M, Tiwari V, Chopra K (2009) Suppression of NF-kβ signaling pathway by tocotrienol can prevent diabetes associated cognitive deficits. Pharmacol Biochem Behav 92:251–259 doi:10.1016/j.pbb.2008.12.012

45 Hu C, Ding H, Li Y et al (2015) NLRP3 deficiency protects from type 1 diabetes through the regulation of chemotaxis into the pancreatic islets. Proc Natl Acad Sci U S A 112:11318–11323 doi:10.1073/pnas.1513509112

46 Evans JL, Goldfine ID, Maddux BA, Grodsky GM (2003) Are oxidative stress-activated signaling pathways mediators of insulin resistance and beta-cell dysfunction? Diabetes 52:1–8

